# Clinically Relevant Mathematical Model for the BCG-based Treatment Of Type 1 Diabetes

**DOI:** 10.1101/2021.09.02.458659

**Authors:** Teddy Lazebnik, Svetlana Bunimovich-Mendrazitsky, Alex Kiselyov

## Abstract

This work introduces a model of immunotherapy treatment, namely the Bacillus Calmette-Guerin (BCG) vaccine, of type 1 diabetes (T1D). The model takes into consideration a clinically relevant interaction network between multiple immune cells and compartments. A set of ordinary differential equations (ODEs) is introduced to capture the connectivity between these variables and clinical presentation of the disease. Four subsets of the T1D patients and healthy controls that exhibit normal and high-level glucose consumption are evaluated. The results that obtained for mice, suggest that BCG treatment of the T1D patients that follow healthy eating habits normalizes glucose to levels observed in non-diabetic controls. Furthermore, glucose consumption profoundly influences disease progression. The stable equilibrium state with constant glucose levels is not attainable without repeated BCG treatment. This outcome suggests that immunotherapy may modulate molecular and cellular manifestations of the disease but it does not eliminate T1D. Of note, our data indicate that the BCG immunotherapy treatment may benefit healthy controls on a high-glucose diet. One may speculate the preventive BCG treatment to provide long-term health benefits in this specific cohort.

**Author summary:** We proposed a clinically relevant mathematical model of Bacillus Calmette-Guerin (BCG) based immunotherapy for type 1 diabetes (T1D) treatment. The model shows that BCG treatment is able to delay the T1D effects and to provide long-term health benefits while it may modulate molecular and cellular manifestations of the disease but it does not eliminate T1D. The main advantage of the proposed model is the ability to personalize the treatment protocol according to the patient’s metabolism and diet.

## Introduction and Related Work

Type 1 diabetes (T1D) is one of the most common chronic diseases of childhood, however clinical symptoms of the disease may occur later in life [1]. In general, the median age for the disease ranges between six years old and puberty [2]. In the past two decades, T1D became a pandemic [3]. In the U.S. alone the number of adult T1D patients was estimated to be 1.25 million (0.5%) in 2017 [4]. Similar statistics in Germany and Norway suggests 2.6% and 3.3% incidence increase, respectively [5, 6].

According to the American diabetes association, diagnosis of diabetes includes a) fasting blood glucose higher than 7 mmol/L, b) any blood glucose of 11.1 mmol/L or higher with symptoms of hyperglycemia, or c) an abnormal 2 hours oral glucose-tolerance test [7]. T1D is believed to be caused by the immune-mediated destruction of insulin-producing pancreatic *β*-cells [8] as evidenced by the presence of a chronic inflammatory infiltrate that affects pancreatic islets [9]. The main treatment protocol for T1D patients in the last few decades includes exogenous insulin replacement therapy [3]. However, this regimen frequently fails to provide metabolic regulation of multiple T1D-related complications that include cardiovascular disorders, neuropathy, and hypoglycemia [10]. Insulin administration *via* pump [11] is associated with imbalanced therapeutic effect, infections, and pump malfunction.

Recent clinical data demonstrate that T1D patients exhibit small-to-none number of viable *β*-cells. Notably, regeneration of *β*-cells in infants and young children could still be detected [12, 13]. This observation implies that both controlled and well-balanced approach to modulating the immune system in T1D patients may improve the production of *β*-cells and reduce circulating glucose levels.

Bacillus Calmette-Guérin (BCG) vaccine is one of the oldest immunotherapies in the clinical practice [14]. BCG treatment of the T1D patients was introduced relatively recently [15]. BCG was originally developed to treat tuberculosis (TB) and early (non-muscle invasive) stages of bladder cancer [16, 17]. A detailed analysis of the literature describes the utility of BCG to a broad spectrum of autoimmune, allergic, and induced adaptive immune conditions [18, 19, 20]. Of particular significance to this work, several instances of BCG vaccines administration in the childhood yielded a dramatic improvement of the T1D progression [21].

Namely, a randomized eight-year-long prospective examination of T1D patients with the long-term disease who received two doses of BCG vaccine was reported by Kühtreiber et al. [15]. The authors showed that BCG furnished a robust and lasting control of the blood sugar levels [15]. Furthermore, the authors discovered that BCG induced a systemic shift in glucose metabolism from oxidative phosphorylation to aerobic glycolysis [22, 23].

Multiple molecular and cellular variables affect the onset of T1D. We reasoned that a clinically relevant mathematical model that recaptures these parameters, as well as the reported BCG-induced clinical observations, may offer a practical insight into immunotherapy of the disease [24, 25]. Earlier, Maree and Kublik [24] introduced an algorithm that describes the interaction between *β*-cells, T-cells, resting macrophages, active macrophages, and immune system-related cells. The authors analyzed the *β*-cells free equilibria which affect T1D. However, the model did not take into consideration the decreased glucose intake, one of the main factor associated with T1D [3]. In our view, this deficiency does not provide insight into the effect of environmental factors, namely dietary glucose. Subsequently, Wu [26] developed an extended strategy that accounts for glucose levels and more detailed cell-cell interactions. The author utilized a multi-compartment model to include the spleen and bloodstream compartments to capture 24 cell types and processes [26]. Whereas the respective single-compartment model reflects data for both healthy and NOD mice, the analysis of the multi-compartment model was not performed due to its size and complexity [26].

Considering the above, we attempted to further streamline the existing mathematical model of the immunotherapy-based treatment of T1D. In this work, clinical markers of T1D including molecular and cellular variables were re-evaluated and the pulsed BCG treatment dynamics introduced by Faustman et al. [27] was applied to the single-compartment model in order to arrive at both relevant and practical outcomes for a mouse model. Our results and presentation are organized as follows:

1. Section 2 provides biological background of T1D and BCG-related treatment used for the mathematical model;
2. Section 3 offers the analysis of the proposed model to yield a set of clinically relevant equilibria states;
3. Section 4 deals with numerical analysis of the model using healthy subjects, T1D patients and corresponding data from the NOD mouse model of T1D. It further defines a baseline behavior followed by the sensitivity analysis of metabolism- and treatment-related parameters including optimal treatment regimen.
4. Section 5 discusses the results including the advantages and limitations of the proposed model, including the personalizing of treatment according to glucose consumption diet, in addition to potential future directions.

## Model Definition

The suggested model is based on the clinically validated pulsed administration of BCG to increase the amount of *β*-cells in the body to prevent T1D [27]. The model describes diverse interactions between BCG and the immune system to arrive at the expected outcome of the treatment on the T-cell and *β*-cell counts.

### Biological background

In T1D or insulin-dependent diabetes, the pancreas produces little or no insulin, a hormone that facilitates glucose to enter cells in order to produce energy. The autoimmune destruction of pancreatic beta cells is one of the key hallmarks of T1D. Glucose metabolism is mediated by multiple types of cells that interact and are controlled by intrinsic and extrinsic factors. Specific cell types include *β*-cells, T-cells, resting and activated macrophages and dendritic cells [28]. The depleted pool of beta cells triggers insulin deficiency and additional pathophysiology including life-threatening hypoglycemia, ketoacidosis, cardiovascular disease, retinopathy, diabetic renal disease, and neuropathy. In a healthy organism, the basic population of *β*-cells is supplemented with additional cells as a response to the increased glucose levels after meals [29].

Once the resulting pool of *β*-cells metabolizes available glucose, they get inactivated during insulin generation [28] and their interaction with Th-cells [30]. This interaction induces production of dendritic cells [28, 29] that in turn prompt resting macrophages to process intracellular antigenic peptides and to eliminate glucose from the circulation [29]. Notably, active macrophages produce IL-1 and TNF to recruit resting macrophages. Once this step is completed, active macrophages get deactivated and return to their resting state [28, 29]. Importantly, *β*-cell, T-cell, dendritic cell populations are tightly controlled to exhibit exponential decay [28]. It is speculated that as BCG is distributed and eliminated post-injection, it engages the dendritic cells and activated macrophages [31, 32] to result in the elevated *β*-cells concentration and overall balancing of glucose metabolism. A summary of this cellular interaction network is shown on Fig. 1.

**Fig 1.**
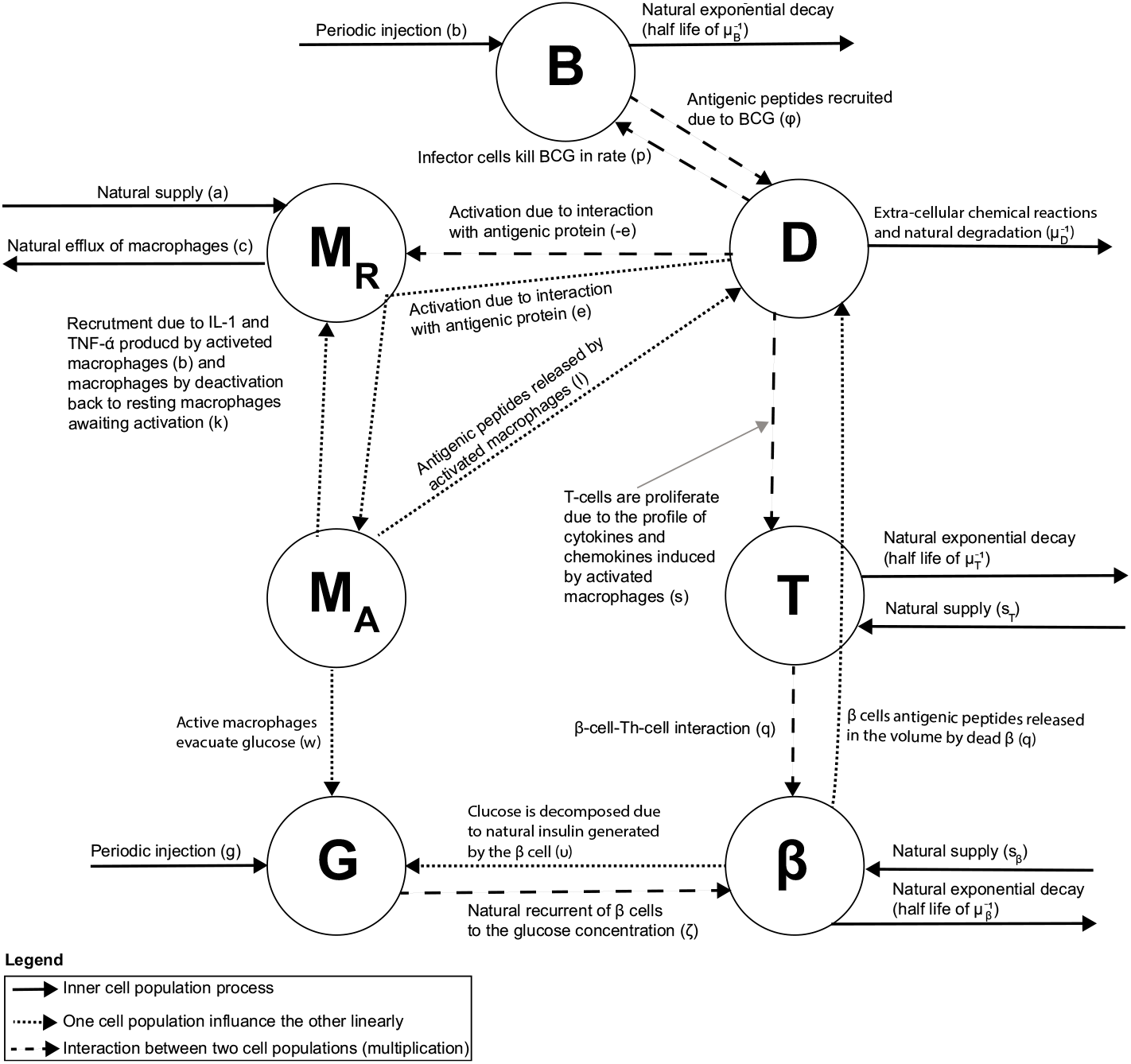
A schematic view of the interaction between the cell populations, divided by interaction type.

Considering the evidence accumulated thus far, significant attention was dedicated to the design and development of immunotherapies to address T1D. Of these agents, the BCG vaccine attracted particular attention primarily due to an array of immune and induced adaptive immune responses exemplified by its clinical performance in multiple sclerosis [33] and non-muscle invasive bladder cancer [34]. Notably, the BCG vaccine was reported to activate immune cells and accelerate glycolysis [32]. Bolus injection of the BCG vaccine-induced significant remission in T1D patients vs controls [31].

### Mathematical model

The suggested mathematical model is aimed to be both clinically accurate and feasible to yield practical analytical insight into the effect of immunotherapy on T1D. We selected seven key variables of interest reported in the clinical practice, namely: *B*(*t*) - concentration of BCG in the body; *M_R_*(*t*) - concentration of resting macrophages; *M_A_*(*t*) - concentration of activated macrophages; *D*(*t*) - concentration of dendritic cells which are also operating as activated immune-system cells; *G*(*t*) - concentration of glucose; *T* (*t*) - concentration of autolytic *T* cells; and *β*(*t*) - concentration of *β* cells.

Furthermore, our approach accounts for cell count dynamics resulting from cell-cell interactions as well as pharmacokinetics and pharmacodynamics processes. These processes are captured using the following system of non-linear, ordinary differential equations (ODEs).

Eq. (1), the 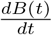 reflects varying levels of free BCG as a function of time. The dynamics is affected by positive and negative processes as follows: i) a BCG installation of concentration/activity *b* > 0 is injected *N* times, every *τ_B_* > 0 time units (which represented using a Dirac delta function [35]). ii) BCG is depleted as a result of natural exponential decay with half life of 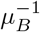. iii) BCG levels decrease proportionally *p* > 0 to the interactions with effector cells including macrophages and dendritic cells [36].

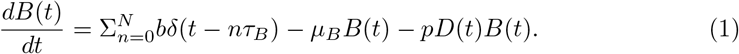

Eq. (2), the 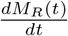 describes the amount of resting macrophages as a function of time. This process is affected by several positive and negative factors. Positive factors are: i) the endogenous supply or recruitment rate of macrophages *a*, ii) recruitment rate *γ* of macrophages due to IL-1 and TNF-*α* produced by activated macrophages and iii) rate of addition of macrophages by deactivation back to resting macrophages awaiting activation. Negative factors include i) rate of decrease in the macrophage population *c* due to the natural efflux of macrophages [24] and ii) rate *e* of macrophages activation due to their interaction with dendritic cells; this factor is proportional to the antigenic peptide and macrophage’s population size.

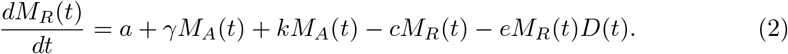

Eq. (3), 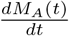 reflects the amount of activated macrophages as a function of time. This variable is affected by the interaction between macrophages and dendritic cells at the rate *e* to increase the number of activated macrophages. Processing of the intracellular antigenic peptides by activated macrophages affords the opposite effect on the macrophage population to lead to resting macrophages at the rate *k*.

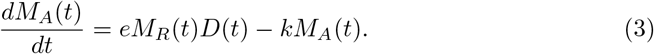

Eq. (4), 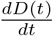 describes levels of dendritic cells as a function of time. The variable is affected by i) *β* cell antigenic peptides released by *β* cells as a result of cell–cell interactions with the autolytic Th-lymphocytes [37]; ii) antigenic peptides produced by the BCG infected cells at the rate *ϕ*; iii) dendritic cells that are cleared at the rate *μ_D_*.

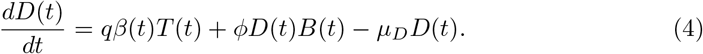

Eq. (5), 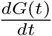 introduces the amount of glucose as a function of time. The variable is affected by i) dietary glucose in the amount of *g* > 0 that is introduced every *τ_G_* time units (represented using a Dirac delta function [35]); ii) active macrophages that eliminate glucose at the rate *w*; iii) glucose consumption due to the natural insulin generated by the *β* cell and released by degrading *β* cells at the rate *υ* [38].

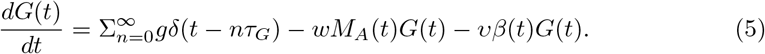

Eq. (6), 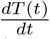 defines levels of autolytic T cells as a function of time. This variable is affected by i) the natural supply or the replacement rate *s_T_* of autolytic T cells; ii) proliferation rate *s* due to the immune cells-induced release of cytokines and chemokines iii) the T cell population decrease due to lack of stimulation at the rate of *μ_T_*.

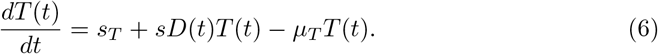

Eq. (7), 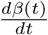 describes levels of *β* cells as a function of time. This variable is influenced by i) the natural supply or replacement rate *s_B_* of *β* of beta cells; ii) glucose-dependent natural production of *β* cells at the rate *ζ*; iii) *β* cell count decrease due to *β* cell-Th-cell interaction[39]; iv) *β* cells count decrease due to the natural attrition at the rate *μ_B_*; v) *β* cells count decrease in a rate *ξ* due to generation of insulin in order to process the glucose.

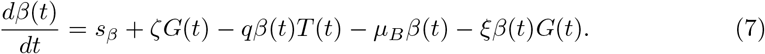

Considering the above, we arrive at the following system of non-linear, first-order, fully-depended ODEs:

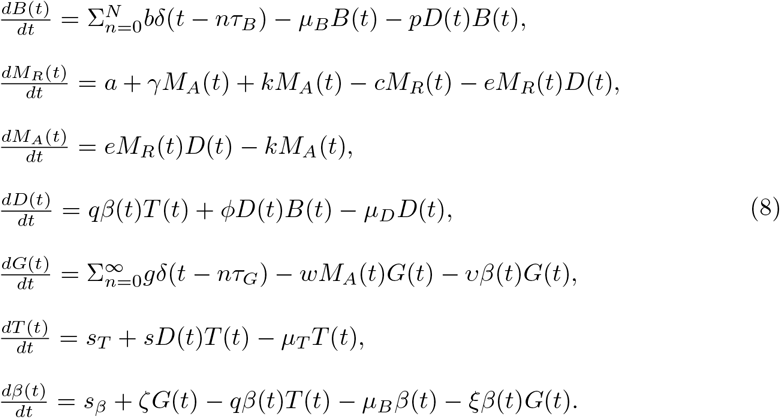

Average parameter values for the model are summarized in Table. 1, for the case of a mouse.

**Table 1.** Parameter description, their average values, and sources for the model.

| Parameter | Description | Average value | Source |
| --- | --- | --- | --- |
| $b$ | The amount of BCG [1]. | $10^4$ | Estimated |
| $N$ | The number of BCG treatment [1]. | 2 | [15] |
| $\tau_B$ | Number of days between any two injections [t]. | 28 | [15] |
| $\mu_B$ | The half-life of the natural decay of BCG expressed in days $[t^{-1}]$. | 0.1 | [40] |
| $p$ | Fraction of BCG decay due to interactions with effector cells expressed in days $[cell^{-1} \cdot t^{-1}]$. | $3 \cdot 10^{-2}$ | [40] |
| $a$ | The natural supply rate of macrophages expressed in days $[mm^{-3}t^{-1}]$. | 50 | [24] |
| $\gamma$ | The recruitment rate of macrophages due to IL-1 and TNF-α produced by activated macrophages expressed in days $[t^{-1}]$. | 0.3 | [24] |
| $k$ | The rate active macrophages are deactivated and becoming resting macrophages expressed in days $[t^{-1}]$. | 0.2 | [24] |
| $c$ | The rate the macrophage population naturally decrease expressed in days $[mm^{-3}t^{-1}]$. | 0.1 | [24] |
| $e$ | The rate of activation for macrophages due to the interaction with dendritic cells expressed in days $[t^{-1}]$. | $5 \cdot 10^{-4}$ | [24] |
| $q$ | The rate of release for antigenic peptides by decaying $\beta$ cells due to cell-cell interactions between the $\beta$ cells and the autolytic Th-lymphocytes expressed in days $[mm^{-3}t^{-1}]$. | $2 \cdot 10^{-6}$ | [24] |
| $\phi$ | The rate of recruitment for antigenic peptides due to the BCG infection of cells expressed in days $[t^{-1}]$. | $3 \cdot 10^{-7}$ | [40] |
| $\mu_D$ | The rate of clearance for dendritic cells via extra-cellular chemical reactions and natural degradation expressed in days $[t^{-1}]$. | 0.5 | [41] |
| $g$ | The average amount of consumed glucose per day in milligrams $[g \cdot t^{-1}]$. | 30 | [42] |
| $\tau_G$ | Glucose treatment intervals expressed in days [t]. | 0.34 | Estimated |
| $w$ | The rate of glucose clearance by active macrophages expressed in days $[t^{-1}]$. | 0.1 | Estimated |
| $v$ | The rate of glucose consumption due to the endogenous insulin produced by $\beta$ cells expressed in days $[ml \cdot t^{-1}]$. | 0.072 | [26] |
| $s_T$ | The supply or replacement for autolytic T cells expressed in days $[mm^{-3}t^{-1}]$. | 20 | [25] |
| $s_\beta$ | The supply or replacement rate for $\beta$ cells expressed in days $[mm^{-3}t^{-1}]$. | 1000 | [25] |
| $s$ | The rate of proliferation for T cells induced by cytokines and chemokines from activated macrophages expressed in days $[t^{-1}]$. | $5 \cdot 10^{-4}$ | [25] |
| $\mu_T$ | The rate of T cell population decrease due to lack of stimulation expressed in days $[t^{-1}]$. | 50 | [24] |
| $\mu_\beta$ | The half-life of $\beta$ cells due to the natural decay expressed in days $[t^{-1}]$. | 0.5 | [26] |
| $\xi$ | The rate of $\beta$ cells decay due to production of insulin induced by glucose [1]. | 1 | [26] |
| $\zeta$ | The rate of $\beta$ cells regeneration induced by glucose $[mm^{-3}t^{-1}]$. | 10 | [26] |

## Equilibria and stability analysis

Equilibrium state presumes that the dynamic system does not change without external stimuli. Stable equilibria are not affected by minor factors which can be taken out of consideration when modeling complex biological systems. In our proposed model (see Section), the equilibria of interest presume the dendritic cell population is equal or close to zero. This is owing to either weak immunologic response caused by the disease burden or due to the weakened response by the immune system to the injected BCG. Therefore, we assume *D*(*t*) = 0 which leads to an equilibrium:

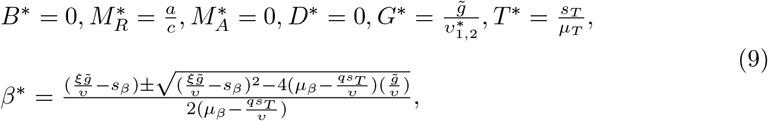

where 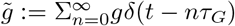 such that *qs_T_ = μ_β_μ_T_*, *υμ*_*β*_ *qs_T_*, and 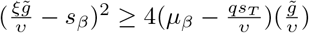. Indeed, we obtain *B\** = 0 as expected. In addition, 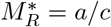 which is the normal rate of *M_A_* cells due to the recurrent and natural decay of the *M_R_* cells population indicative of the non-compromised immune response due to the treatment. Furthermore, 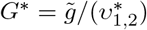 which implies that the glucose levels are periodic and exhibit a first-order influence on the amount of *β*-cells.

For the system of Eq. (8), we evaluated the Jacobian *J* :

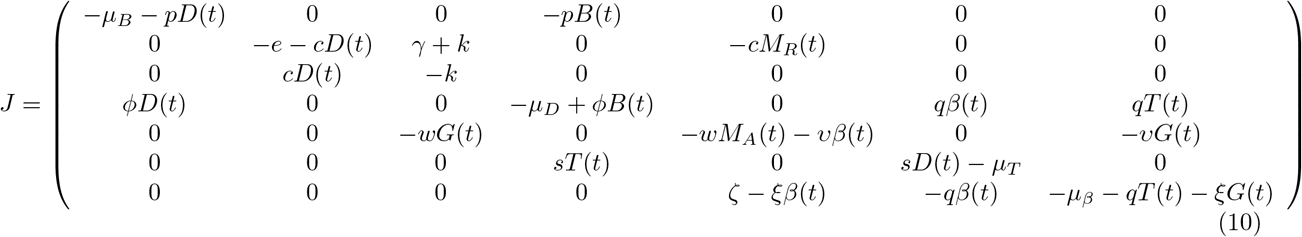

By setting the values from Table 1 and Eq. (9) in Eq. (10), one gets that Eq. (9) is indeed locally stable.

In a physiologically relevant equilibrium a healthy patient is expected to maintain the glucose levels within ‘normal’ manageable margins *G*(*t*) ∈ [*l_b_, u_b_*]. We define the *healthy* equilibrium to be the steady state in which the glucose levels are in the middle of this range (e.g., *G\** = *h* = (*l_b_ + u_b_*)*/*2) and no BCG is administered (*B\** = 0). In this case, the equilibrium looks as follows:

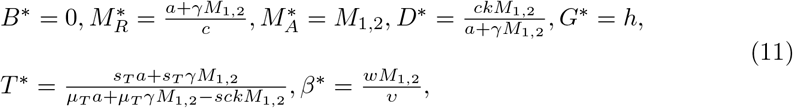

such that

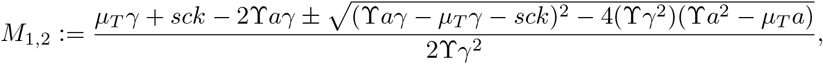

where 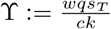 and (ϒ*aγ* − *μ_T_ γ* − *sck*)^2^ ≥ 4(ϒ*γ*^2^)(ϒ*a*^2^ − *μ_T_ a*). This equilibrium is not biologically stable as 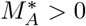 which is an indication for an activated immune response. This outcome means the individual is affected by the disease. Therefore, in order to evaluate the case where 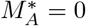, we get that

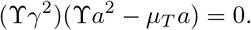

Therefore,

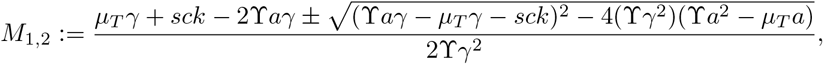

As *w, q, s_t_, γ, a, μ_T_* > 0 by definition, the equilibrium takes the form:

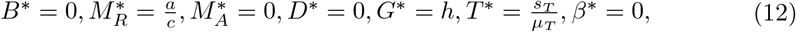

and due to the fact that *β\** = 0 the glucose levels raise to lead to the unstable equilibrium as well as to unrealistic clinical scenario. As a result, the equilibrium in Eq. (11) is unstable.

## Numerical Simulations

We solve the model (Eq. (8)) numerically using the *ode15s* function in Matlab (version 2020b) [43, 44], where the model’s initial conditions are as follows [15]:

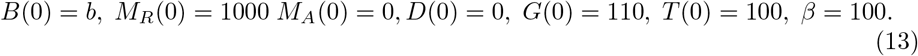

The model’s parameter values are taken from Table 1 and represent the average values for a mouse. We evaluate the baseline dynamics of the model using four types of individuals. The first type is the non-T1D patient which eats healthy and consumes approx. 30 milligrams of glucose per day. The second type is the non-T1D patient eats over 90 milligrams of glucose per day. The third type is the T1D patient eats healthy. Finally, the T1D patient shows unhealthy glucose consumption habits. The results are shown in Figs. 2a-2d, where the x-axis is the post-BCG treatment time and the y-axis is reflective of the cell population size. As expected, the model shows that the baseline treatment protocol defined by the values in Table 1, does not affect healthy individuals (Figs. 2a-2b. Importantly, the protocol delays T1D markers (e.g., *G* > 110 for a day on average) in T1D affected cohort on a healthy diet (Fig. 2c. The baseline treatment protocol is not effective in T1D patients on high glucose diet.

**Fig 2.**
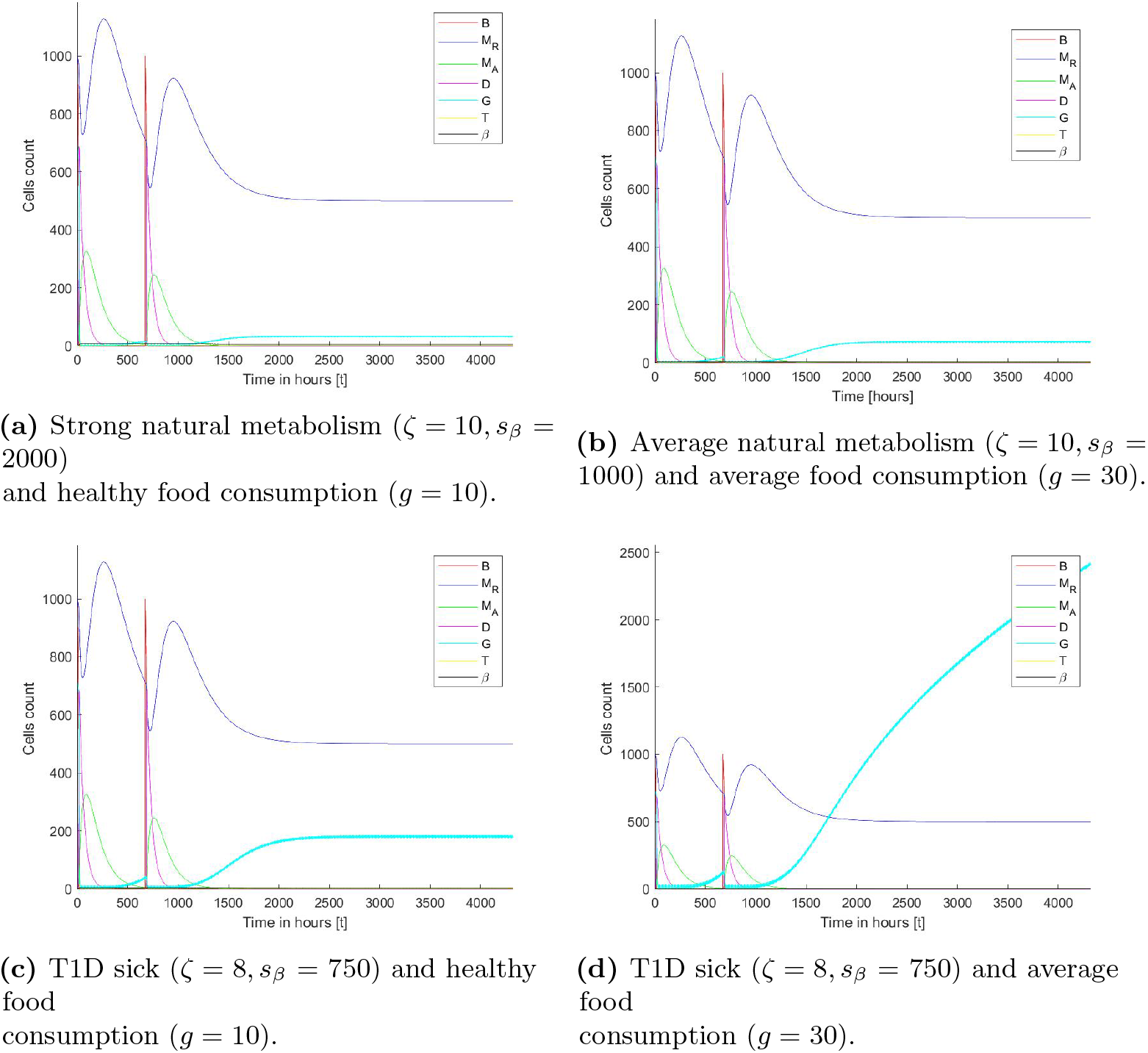
Dynamics of T1D-relevant cell markers in 4 distinct cohorts including healthy subjects and T1D patients as a function of time. The parameters are taken from Table 1 and the initial condition from Eq (13).

Our data suggest that there is a relationship between the patient’s dietary habits, such as daily glucose intake (*g*) and genetic factors, such as metabolism and *β* cell production due to glucose stimulus (*s_β_*). Sensitivity analysis *g* ∈ [0, 50], *s_β_* ∈ [800, 1200] shows the effect of both parameters on glucose levels three months after the last BCG treatment (Fig. 3).

**Fig 3.**
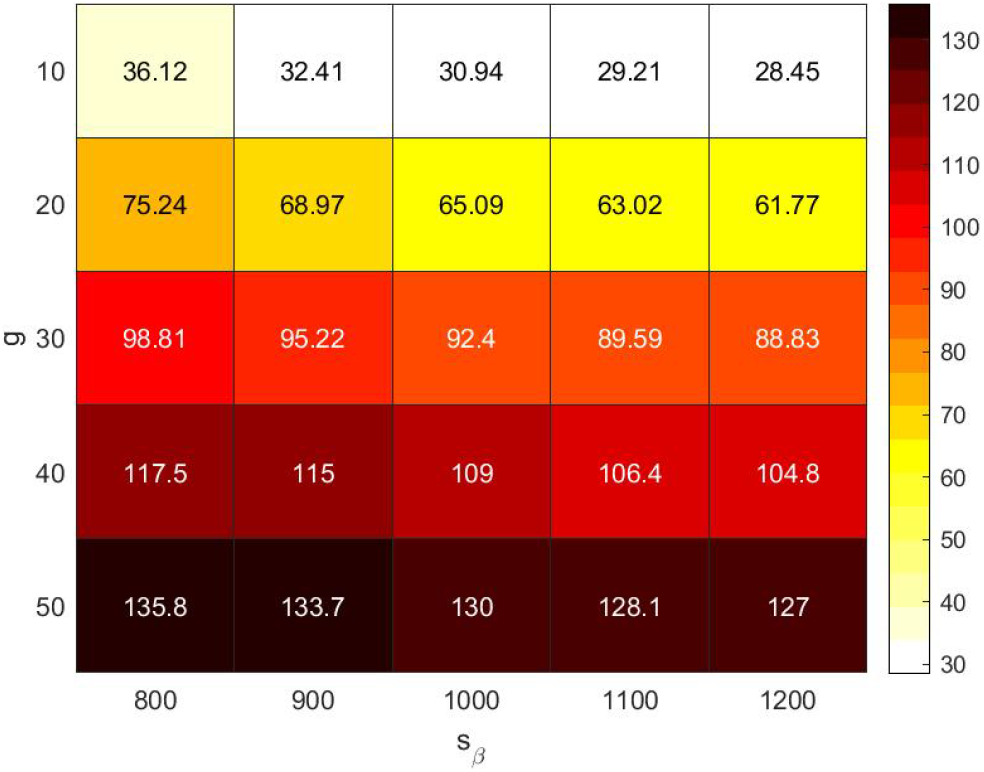
Sensitivity analysis of average glucose consumption per meal *g* (*τ_g_* = 8 hours) and recruitment of *β* cells *s_β_* indicative of the baseline metabolism on the daily average glucose levels three month after the last BCG treatment (*N* = 2, *b* = 1000).

The values obtained in the sensitivity analysis are fitted using the least mean square (LMS) method [45]. The selected family function for the surface approximation is as follows:

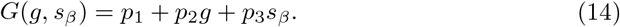

A linear approximation is chosen to balance between the accuracy vs the sampled data on the one hand and model’s simplicity and interoperability on the other hand [46], which results in:

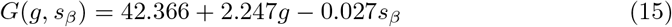

to afford a coefficient of determination *R*^2^ = 0.749. Therefore, the influence of *g* is estimated to be 10^2^ times larger than the influence of *s_β_*.

Because the eating habits and physiology are unique to each patient, the optimal treatment is expected to differ between patients. In order to find the most relevant patient-specific regimen, one can further analyze the image of the model (e.g., Eq. (8)) by solving the ODE and to produce a function *T* (*b, τ_b_, N*) → {0, 1} when the source space contained in ℝ^3^ and the image space is exactly {0, 1} [47]. The parameters (*b, τ_b_, N*) are used as the source space since they define the BCG-based treatment protocol. The other parameters are patient-dependent and can not be modified as a part of the treatment. The results of the analysis for each individual patient are summarized on Fig. 2. The green (circle) and red (star) dots represent successful and unsuccessful treatments, respectively, such that the evaluation performed six month from the last BCG treatment (which differ as a function of *N*), in Fig. 4.

**Fig 4.**
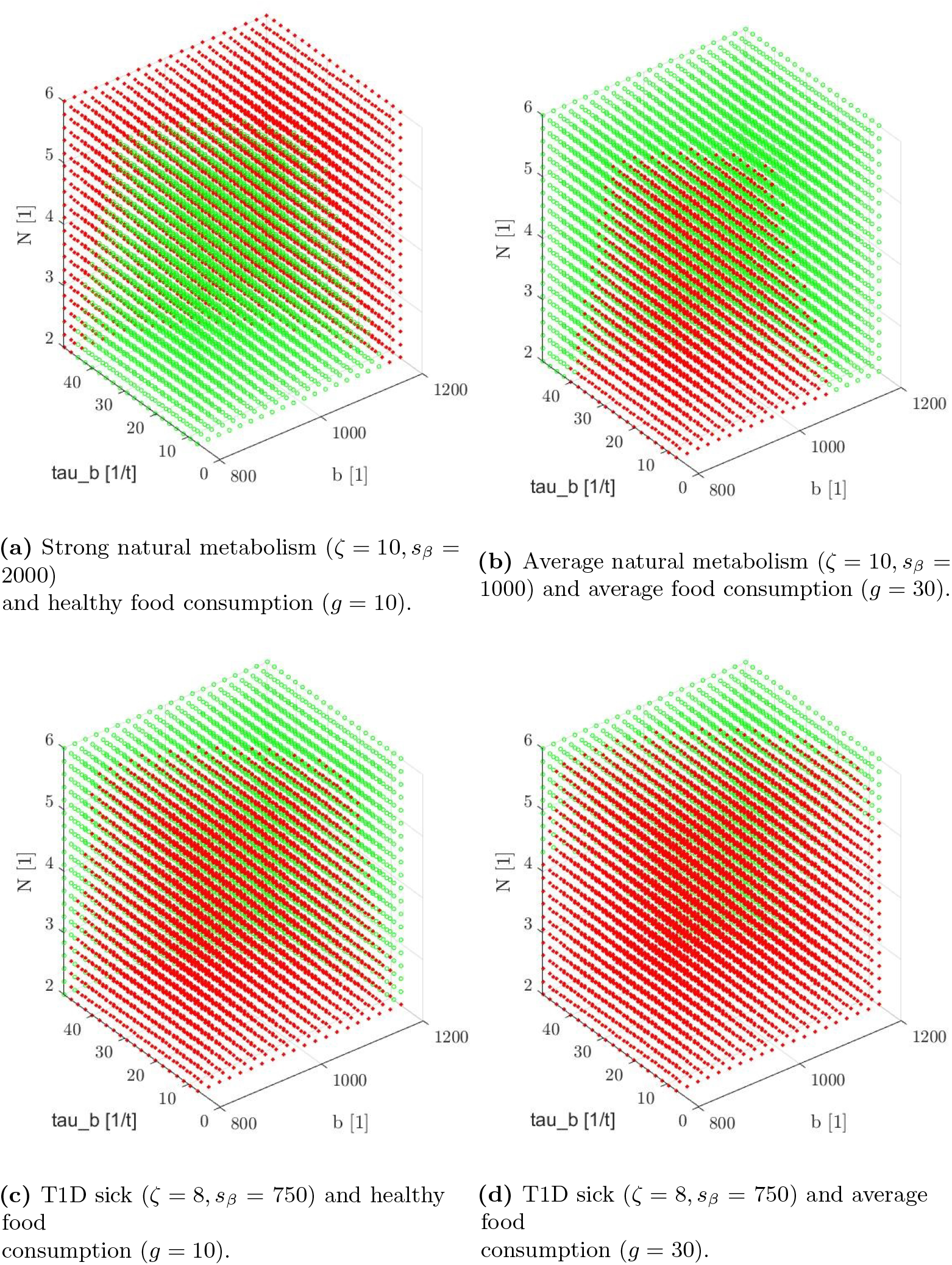
Discrete sampling of the model’s image space for the domain (*b* ∈ [800, 1200], *τ_b_* ∈ [10, 50], *N* ∈ [2, 3, 4, 5, 6]). Green (circle) pixels represent successful treatment and red (star) pixels represent unsuccessful treatment. The parameters taken from Table 1.

For each patient, we find the set of border pixels (neighboring pixels that show both colors) indicative of the separation between successful and unsuccessful treatments. The set of border pixels defining a surface can be approximated using the least mean square (LMS) method [45]. The family function for the surface approximation is selected as follows:

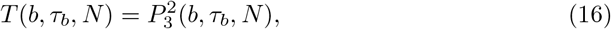

where 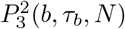 is a second order three-parameter polynomial. This function is selected manually (after testing multiple functions) to balance between the coefficient of determination and simplicity of the model. The results of the approximation are as follows:

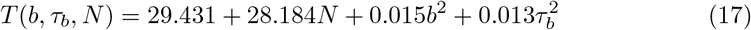

to yield the coefficient of determination *R*^2^ = 0.920 for Fig. 4a. Similarly, for Fig. 4b:

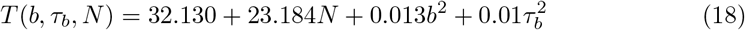

to result in a coefficient of determination *R*^2^ = 0.912. Similarly, for Fig. 4c:

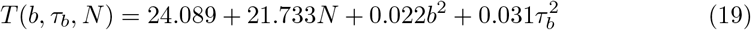

to afford a coefficient of determination *R*^2^ = 0.897. Similarly, for Fig. 4d:

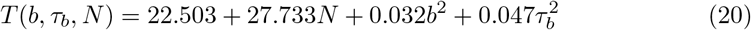

to produce a coefficient of determination *R*^2^ = 0.874.

## Conclusion

In this work we introduced a model of immunotherapy treatment of type 1 diabetes (T1D) using BCG. As multiple cell types and organs are involved in controlling intricate metabolic balance both in healthy subjects and T1D patients, we introduced a set of specific variables (markers) immediately relevant to the published clinical data on the disease. Furthermore, a system of ODEs reflecting the molecular, cellular, pharmacokinetic, and pharmacodynamic connectivity between these variables was introduced to capture the clinical presentation of the disease in four subsets of the T1D patients and controls.

In our view, the model (Fig. 2) accurately predicts the observed fundamental importance of an appropriate diet regimen. Namely, BCG treatment of the T1D patients that follow healthy eating habits normalizes glucose to the levels observed in non-diabetic controls. This outcome indicates the initial validation of the introduced model. In the context of the BCG therapy, our data propose that glucose consumption exert more influence (est. 10^2^ difference) on the disease progression than metabolism, especially when evaluated over a clinically relevant treatment time (Fig. 3).

Figs. 4c and 4d illustrates the profound effect of glucose on the outcome of the T1D. Namely, T1D patients featured in Fig. 4d are expected to perform much worse compared to patients shown on Fig. 4c even following the insulin treatment. Intriguingly, data analysis summarized on Fig. 4d indicates that the BCG treatment may still benefit healthy controls on a high-glucose diet. One may speculate the preventive BCG treatment to provide long-term health benefits in this specific cohort.

Eqs. (17–20) represent the border function between the successful and unsuccessful treatment protocols, differing in the amount of BCG injected, the rate and number of the injections.

We conclude that it is possible to find the optimal treatment protocol by setting a specific desired value for properties comprising the protocol to recommend a personalized treatment that corresponds to the patient’s glucose intake.

Moreover, the equilibrium described by Eq. (9) suggests a long-term system stability. This outcome is clinically relevant as it indicates that the BCG treatment may have a defined longitudinal effect and should be repeated as needed. Eq. (11) corroborates this conclusion to imply that a biologically stable state of the system with stable glucose levels is not attainable without repeating the BCG treatment. In other words, the BCG treatment seems to modulate molecular and cellular manifestations of the disease but does not eliminate T1D.

In our view, the proposed system of parameters and equations introduces both accurate and practical clinical models for the immunotherapy-based treatment of T1D. Our model further proposes that additional interventional modalities, for example mechanistically different immunotherapy including vaccines or engineered cells may enhance both targeted and symptomatic treatment outcomes.

